# DeepDBP: Deep Neural Networks for Identification of DNA-binding Proteins

**DOI:** 10.1101/829432

**Authors:** Shadman Shadab, Md Tawab Alam Khan, Nazia Afrin Neezi, Sheikh Adilina, Swakkhar Shatabda

**Affiliations:** Department of Computer Science and Engineering, United International University, Plot-2, United City, Madani Avenue, Badda, Dhaka-1212, Bangladesh

**Keywords:** DNA-Binding proteins, Deep Learning, CNN, Classification algorithm, Feature selection

## Abstract

DNA-Binding proteins (DBP) are associated with many cellular level functions which includes but not limited to body’s defense mechanism and oxygen transportation. They bind DNAs and interact with them. In the past DBPs were identified using experimental lab based methods. However, in the recent years researchers are using supervised learning to identify DBPs solely from protein sequences. In this paper, we apply deep learning methods to identify DBPs. We have proposed two different deep learning based methods for identifying DBPs: DeepDBP-ANN and DeepDBP-CNN. DeepDBP-ANN uses a generated set of features trained on traditional neural network and DeepDBP-CNN uses a pre-learned embedding and Convolutional Neural Network. Both of our proposed methods were able to produce state-of-the-art results when tested on standard benchmark datasets.DeepDBP-ANN had a train accuracy of 99.02% and test accuracy of 82.80%.And DeepDBP-CNN though had train accuracy of 94.32%, it excelled at identifying test instances with 84.31% accuracy. All methods are available codes and methods are available for use at: https://github.com/antorkhan/DNABinding.

## 1 Introduction

DNA is the blueprint for the cell. It contains all the information and instruction that codes for the development and function of living things. But DNA does not do it by itself. There are thousands of DNA-binding proteins that help modulate DNA’s functions. DNA-binding proteins have an indispensable role in major cellular processes. DNA replication and recombination are the two major functions of DNA Binding proteins. However identifying the proteins that bind in the major groove is one of the challenging task.

Having been linked to several cellular functions, it is highly crucial to identify DNA binding proteins. Over the past few years, several traditional machine learning methods have been applied to classify DBPs. Nowadays Machine Learning (ML) algorithms are very effective as computational method to recognize DBPs. Over the past few decades the traditional machine learning methods have proved to be cheaper, faster [1] and more capable to deal with the sudden outburst of data compared to any other methods and hence have been extensively used in many different papers.

The sequence based predictors got the most attention of researchers to improve the performance of identifying the DNA binding proteins as that do not need the information of protein sequence structure. Feature representation and classification algorithms are the most important components to perform a ML-based method for identification of DNA binding protein. Numerical feature representation is the best way of representing of protein sample. There are mainly two categories predictor for feature representation based on ML: i) structure-based predictors and ii) sequence-based on predictors.

The breakthrough idea of pseudo amino acid composition [2] or PseAAC [3, 4] was proposed by Chou and from then on has been used in countless number of papers [5, 6, 7, 8, 9, 10, 11, 12, 13, 14, 15, 16, 17, 18, 19]. The concept of Pse-AAC have been directly applied in several state of the art models like DNABinder [20], BLAST [21], PseKNC(Pseudo Ktuple Nucleotide Composition) [17], etc. Even better results were obtained when machine learning algorithms like Random Forest, Support Vector Machine, etc. were incorporated to the model. DNA-Prot [22] was initially trained using Random Forest(RF) classifier and was later named iDNA-Prot in [23] after the addition of Grey Model. RF was also used in the training of the Local-DPP [24] model. Support Vector Machine(SVM) was used in iDNAPro-PseAAC [1] to improves predicting capability. The model was later made faster with the help of dimension reduction and was renamed to iDNA-Prot|dis [25]. Both the SVM and RF classifiers were used in Kmer1 + ACC [26]. Feature selection was carried out using RF classifier in both DBPred [27] and DPP-PseAAC [28].

Biological information like structural and evolutionary information were added to obtain better results in HMMBinder [29] and iDNAProt-ES [30]. A similar approach was applied, with the combination of RF and SVM, in iDNAPro-PseAAC [31]. Moreover, a bunch of web-servers and open source tools like Pse-in-One [32] and Pse-in-One2.0 [33], PseAACBuilder [34] and PseAAC-General [35] were also made public for the aid of scientists across the globe.

Due to the development of next generation sequencing (NGS), also known as high-throughput sequencing, it is now much easier to sequence DNA and RNA quickly; as a result the number of new protein sequence has increased. According to The Universal Protein Resource Knowledge-base(Uniprot), protein sequence repository is increasing. Even though the experimental approaches used over the past few years are able to identify DBP’s correctly, they are slowly becoming less effective with the increase in data. Therefore more efficient and time-saving computational methods need to be introduced to manage the increasing protein sequence data to identify DBPs.

In the early 2000s a new approach began to emerge among the researchers. The use of Deep Learning(DL) started to become more popular in the field of Bioinformatics. While the traditional ML approaches failed to process the huge amounts of data efficiently, the DL methods showed tremendous efficiency. DL is a novel approach and was inspired from the neurons in human brain. This approach is able to work with raw data and does not require the features to be extracted prior to processing [36, 37, 38]. It is still an abstract and complicated approach and the network architecture is still a black box to the scientists [39]. In 2015, DeepBind software [40] was created which was able to predict the DNA and RNA sequences using DL. In 2016, an approach with a relative 50% improvement was introduced and was named DanQ [41]. KEGRU [42, 43], introduced in 2018, is a Recurrent Neural Network(RNN) based architecture which uses k-mer embedding combined with a layer of GRU units. Hybrid approaches combining Convolutional(CNN) and Recurrent Neural Networks were used to predict enhancers in DNA. The Biren [44] method took solely the DNA sequence and did the rest of the processing on its own. The concept of DL has been used in several other researches and is still being used till date due to its efficiency and novelty.

The authors of [45] compared three different types of models: RNN, CNN and hyrbid of CNN and RNN. They introduced several new techniques inspired from exising models like deepBind, DanQ, etc. All their models can be found in their online tool called deepRAM. Their experimental results prove that the hybrid models outperform the solo architectures. They named their two best architectures ECBLSTM and ECLSTM both of which were created using k-mer mebedding and hybrid neural network layer. The experiments where all carried out on the ChIP-seq ans CLIP-seq from the ENCODE project [46, 47, 48]. Even though the hybrid models have outstanding performance on RNA-binding proteins, a significant drop in performance can be seen when tested on DNA-binding proteins.

In this paper, we have used two different approaches: DeepDBP-ANN and DeepDBP-CNN. Both of the methods are using deep neural network to solve the DNA binding protein prediction problem. DeepDBP-ANN uses a generated set of features trained on traditional neural network and DeepDBP-CNN uses a pre-learned embedding and Convolutional Neural Network. Both of our proposed methods were able to produce state-of-the-art results when tested on standard benchmark datasets.DeepDBP-ANN had a train accuracy of 99.02% and test accuracy of 82.80%.And DeepDBP-CNN though had train accuracy of 94.32%, it excelled at identifying test instances with 84.31% accuracy.

## 2 Materials and Methods

We have used two distinct approaches for the methodology. The first one includes the 5 standard steps introduced by Chou [49] and summarized by Rahman el al.[50] which follows: i) acquire a standard test and train dataset; ii) represent features in form of a feature vector; iii) develop a classification algorithm; iv) neutrally evaluate the classification algorithm and v) creating public access to the classifier. Our first method DeepDBP-ANN follows this technique and the architecture is shown in Figure 1.

**Figure 1:**
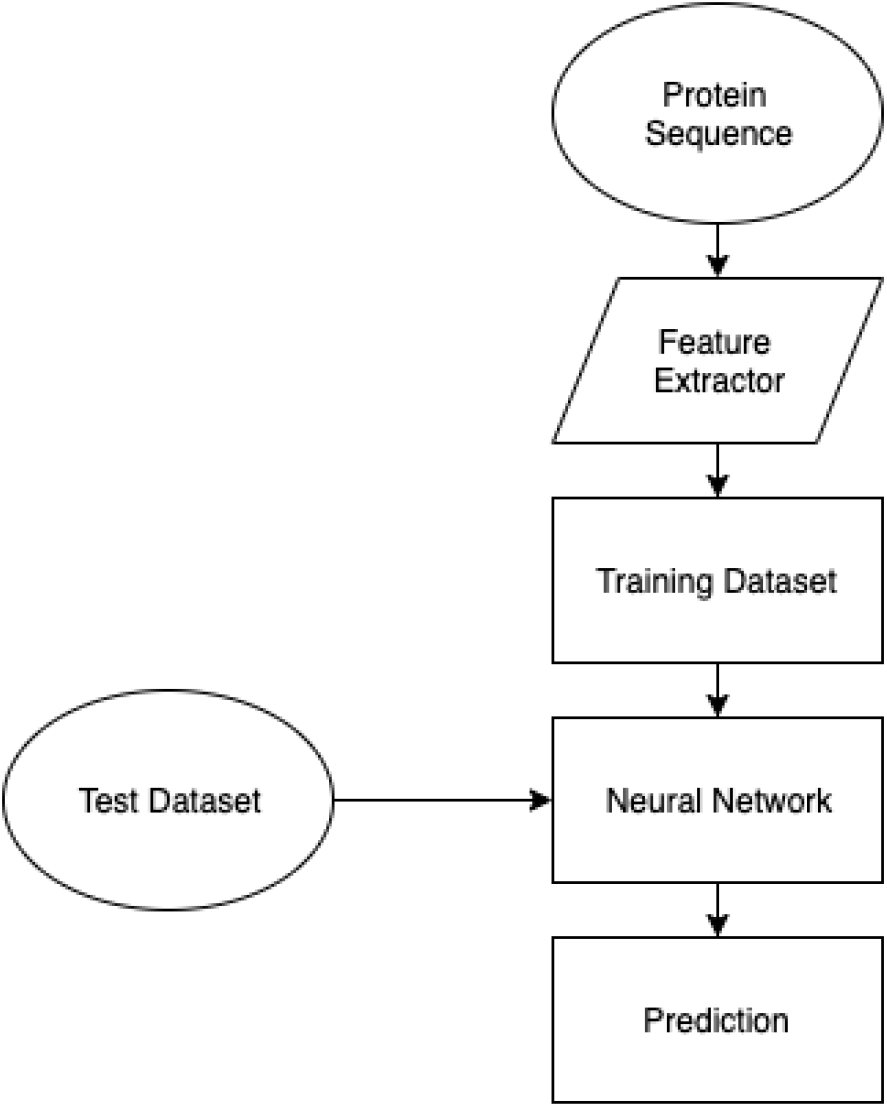
Architecture of the methodology for DeepDBP-ANN.

However this approach has a major drawback, that is it requires various algorithms to extract the features. Those algorithms are often dataset specific. And it needs human interaction to extract the features. Therefore, we came up with another approach which doesn’t require featurized data but rather takes raw amino acid sequence as input. The second approach isn’t dataset specific. The second approach works as follows: we take the model that we’ve discussed in our first section and add a convolutional block to it. Along with the addition on a convolutional block we replace the features that we have used with a set of embedding vectors where each vector represents an *L*-dimensional point. To generate the features from these embedding vectors we create a 2D matrix from a sequence of proteins of length *N*. The size of our final 2D matrix is (*L × N*). On this matrix, we apply repeated convolutions and sub-sampling. Each convolutional is done with a window-size of *L* * 31 and on each layer we apply 128 filters to generate 128 unique feature-maps. These feature-maps are then sub-sampled using max-pooling to shrink the size of the feature-maps. These reduced feature-maps are applied to repeated convolutions and sub-sampling we are left with a *k* feature maps with dimensions of 1 × 1. These feature-maps are treated as *k* features and fed into the neural-network that we developed in the previous step. The architecture is shown in Figure 2

**Figure 2:**
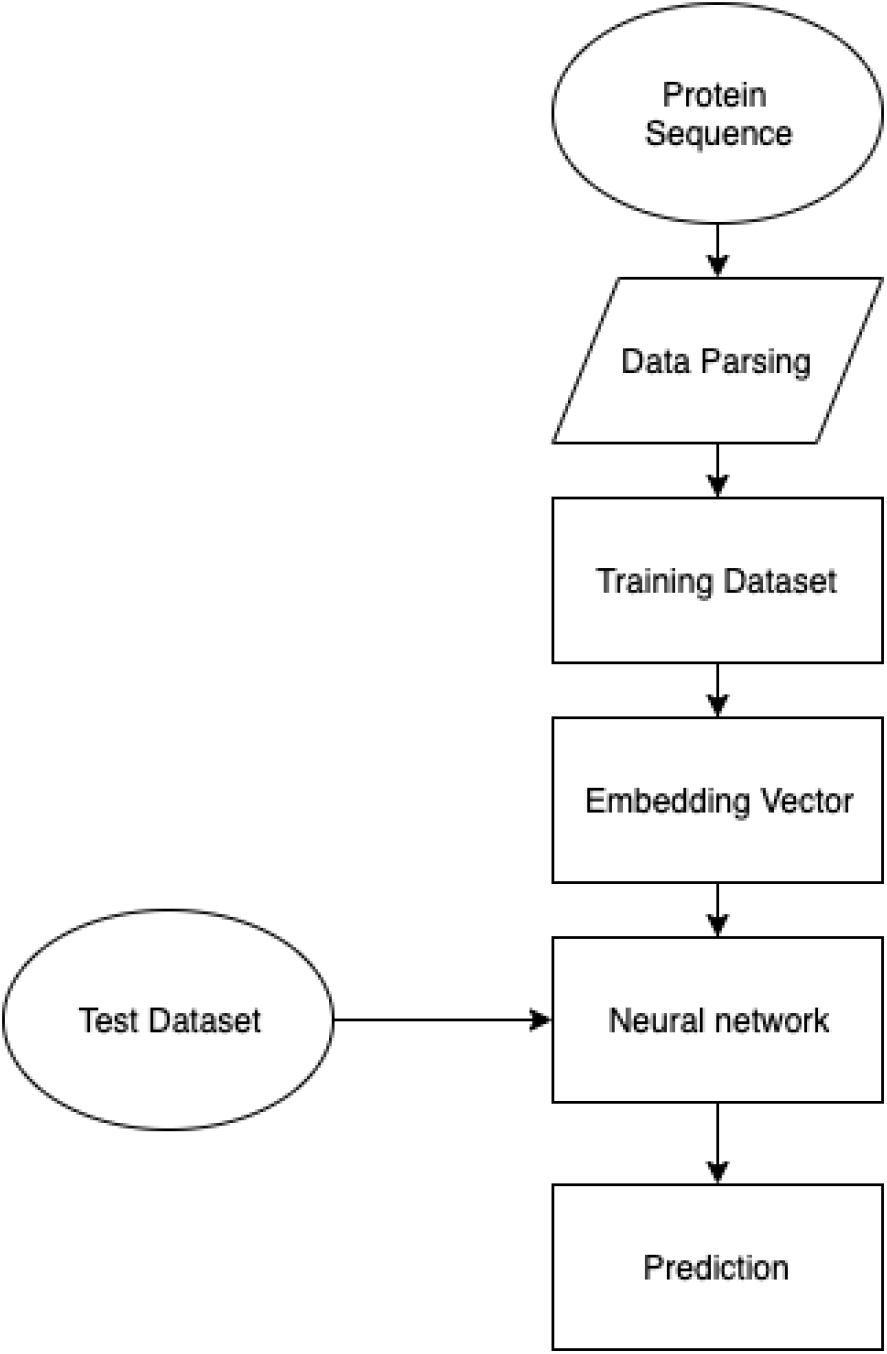
Architecture of the methodology for DeepDBP-CNN.

### 2.1 Benchmark Datasets

For evaluating the predictor, credibility and accuracy of the dataset is crucial. The datasets we used to test our predictor were extracted from Protein Data Bank (PDB: http://www.rcsb.org/pdb/home/dome.do) using certain keywords such as “DNA-binding protein”,”DNA-binding” and so on. Our training dataset PDB1075 was originally extracted by Liu et al.[25]. The PDB1075 dataset contains 525 positive DNA-binding protein sequences and 550 negative sequences. The validation set was compiled by Lou [27],was also extracted from Protein Data Bank contains 93 positive DNA-binding protein sequences and 93 negative ones. Both the datasets have been around for a couple of years,and contains desirable number of protein sequences.

### 2.2 Feature Extraction

As stated earlier we have used two distinct approach for feature extraction and sample representation. In this section, we describe them in two different subsections.

#### 2.2.1 Features for DeepDBP-ANN

Here for DeepDBP-ANN, we used 7 set of features used by Adilina et al. [51] These 7 set of features in total generates 32620 features.

1. **Monograms**:In order to find monograms recurrence of each individual amino acids was determined and then was normalized by the length of the sequence.

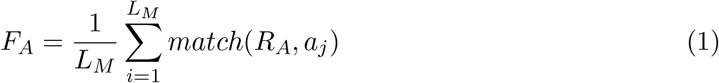

where:

*L_M_* = linear measure of the sequence
*a_j_* = an amino acid from alphabet Σ
*R_i_* = an amino acid at specific position i The function works as following,

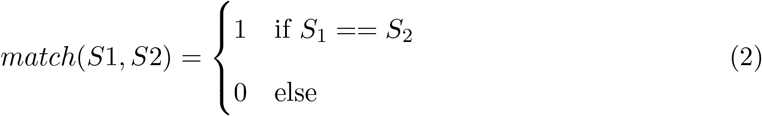 So,There are 20 monograms.
2. **Bigram**: To find the bigrams, recurrence of two successive amino acids was taken into account.Just as the monograms,the frequencies of those bigrams were normalized as well. From the 20 amino acids 400 bigrams are generated.

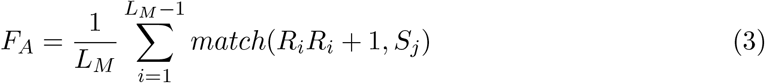

where:

*S_j_* = a di-amino acid string taken from Σ^2^
3. **Trigram**: Trigrams are determined same as the bigrams. Except for two successive amino acids, three are taken into account.

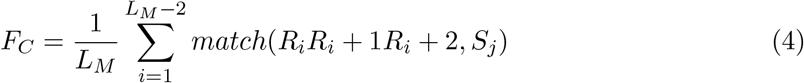

where:

*S_j_* = a tri-amino acid string taken from Σ^3^
4. **Gapped bigram**: Frequency of all possible pairs of amino acids, with a gap of certain length in between, was calculated in Gapped bigrams [52, 53].

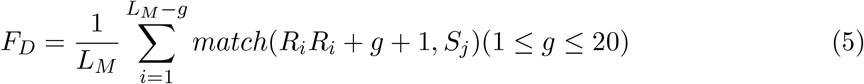

where:

*S_j_* = a di-amino acid string taken from Σ^2^
g = distance between two amino acids in the sequence In this paper, we have used gaps, g = 1, 2,…, 20. Thus the total number of features generated was 80000.
5. **Monogram Percentile Separation**: Here monograms are determined only for partial sequences. On first iteration only 10% of the sequence is taken into account and monograms are determined only for the partial sequence. Then on each iteration we gradually increase the length of the partial sequence by 10% and repeat the whole step until we take 100% of the sequence. We can generate 200 features for each of the protein sequences.

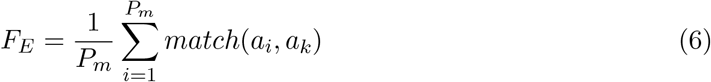

where:

*P_m_* = partial linear measure of sequence
*a_j_* = an amino acid from alphabet Σ
6. **Bigram Percentile Separation**:This process is fundamentally same as monogram percentile separation. Only instead of a individual amino acid, a pair is taken into account. This generates 4000 features in total.

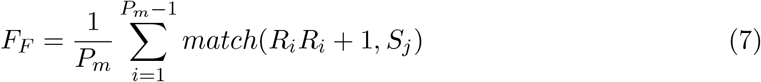

where:

*P_m_* = partial linear measure of sequence
*S_j_* = a di-amino acid string taken from the alphabet Σ^2^
7. **Nearest Neighbor Bigram**:*a_i_* and *a_j_* are considered nearest neighbor bigram[25] if *a_j_* is closest to *a_i_*. Based on this concept, first 30 NNBs are considered to create 12000 features. Just as the previous ones, these values are normalized as well.

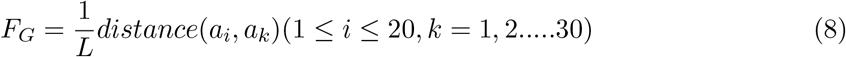

where:

*P_m_* = partial linear measure of sequence
*a_j_* = an amino acid from alphabet Σ

#### 2.2.2 Features for DeepDBP-CNN

We extract the features with the help of a Convolutional Neural Network and an Embedding vector. To deal with sequences of varying length, we pad each sequence to a fixed length by appending a padding token to the end of each sequence. This greatly simplifies our model architecture, as our model can now expect an input of uniform length. We also discuss measures taken such that this does not affect the output of our model. Figure 3 shows the architecture of model of feature extraction mode.

**Figure 3:**
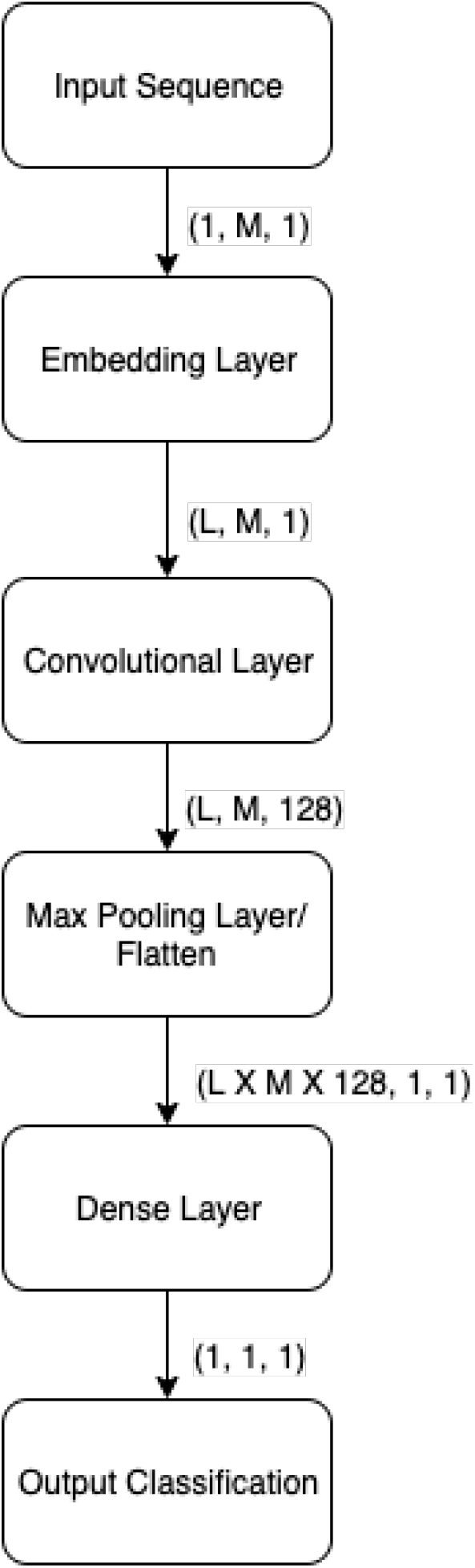
Architecture of Feature Extraction for DeepDBP-CNN.

#### 2.2.3 Embedding Layer

The first layer in our model is a trainable embedding layer. Embedding layers are used to transform discrete inputs to points in vector space, called embedding vectors. Embedding vectors are a staple of natural language processing where they are used to represent words in an L-dimensional space, where L is the length of the vector. The distance relationships among the vectors, are a representation of their relation to one another. In natural language processing they represent the relation among input tokens, which are in general words, in a fixed set of possible words, i.e. a dictionary. For our model we consider each protein to be a discrete input token, and the set of 20 proteins to be our dictionary. We chose to ignore non-protein tokens, such as tokens used to pad our sequences, by representing them as zero vectors in our embedding space. This zero vectors do not have any affect on the outputs of the subsequent layers. The final output of the embedding layers is a uniform matrix of size LxM where M is the length of our input sequence.

**Figure 4:**
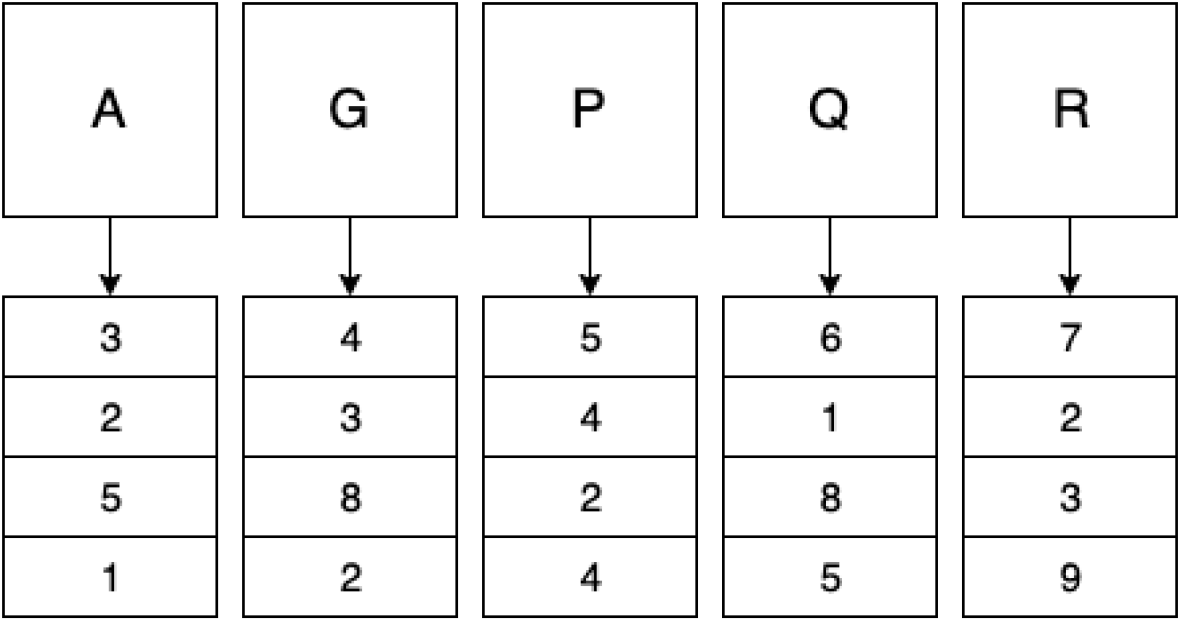
Protein Embedding

#### 2.2.4 Convolutional layer

The next layer in our model is a trainable convolutional layer. The convolutional layer receives uniform matrices generated from our embedding layers, and applies convolutions using 128 trainable filters, each with a window of size Lx31. The result is 128 feature maps, each of the same size. To reduce over-fitting and capturing noise, these feature maps are then sub-sampled using max pooling, with a window size of size 3×3. Finally we flatten the feature maps to 1×1 dimensional matrices, where each matrix represents a feature of the input sequence. These features are then used to train our classification model.

The unique benefit of this approach is that the results of the classification model can be back-propagated to the convolutional and embedding layers, training the layers to extract better features, effectively making our feature extraction trainable.

### 2.3 Classification Algorithms

For the purpose of classification we used deep artificial neural network. The deep learning model was composed of 12 layers excluding input and the output layer. There are four types of layers in our model; dense,activation,batch normalization and dropout. There’s a brief description on each types of the layers below. In a dense layer (also called a fully connected layer) each of the input nodes are connected to each of the output nodes.

#### 2.3.1 Activation Function

An activation node determines output of one or more nodes through a function. We used the Relu as our activation function. [54] Relu functions, shown on equation 9 scales the output in a range of zero to one.

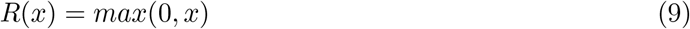

#### 2.3.2 Dropout Layer

Next is the dropout layer. It is an elegant and simple way of dealing with overfitting which has been a challenge for deep neural net for a prolonged period of time. As mentioned by Srivastava et al.[55] dropout means randomly dropping a node along with all its connections from the network. In the process of randomly dropping a node it makes the network less dependent on a single node and thus reducing overfitting.

#### 2.3.3 Batch Normalization Layer

Finally, the batch normalization layer. During the time of writing the document batch normalization was perhaps one of the single most important discoveries that happened in the field of deep learning.Change in distribution of each layer’s input makes training Deep Neural Network difficult. With the change in the parameters of the previous layers, the distribution of the next layer’s input also changes hence making training of deep neural network complex. This decelerates the training by requiring lesser learning rates and cautious parameter initialization, and makes it notoriously tough to train models with saturating non-linearities. Sergey et al.[56] called this problem internal co variate shift, and it can only be solved by normalizing layer inputs.

To summarize, what we normally do is normalize the inputs of a network. So the general assumption was if inputs layers can be benefited by normalization, so should be the hidden layers as well. Batch normalization minimizes the amount by which the hidden unit values deviates. On top of that batch normalization qualifies each of the layers of the network learn by itself independently of other layers. As mentioned by Sergy I. and Christian S. batch normalization has the following qualities among many others.

1. Batch normalization qualifies a network to have higher learning rates. Having the learning rate set to too-high may make the gradient get stuck at local minima or as well as explode or vanish. By normalizing the activations all through the network, batch normalization prevents insignificant change into parameters to being amplified into a large and sub-optimal changes.
2. While using Batch Normalization, a training example is used together with other examples in mini-batches, it no longer produces deterministic values for a given training example. In short it generalized the network and in the process diminish the need of Dropout[55].

#### 2.3.4 Model Architecture

We started our experiment with a simple linear classifier and gradually added more layers as well as nodes on the layers. As seen on our experiment, we are benefited from layers to the architecture, but that is upto a certain point, adding more layers after that point only added overhead to the training process with little or no benefit. It was determined from experiment that 3 layers are the threshold value for number of layers. Adding more layers doesn’t further help our cause. And for the number of nodes on each layer the idea was simple. The more nodes on the layer, the better it performed. So we kept adding more nodes to the layers until we ran out of VRAM on our GPU.

### 2.4 Performance Evaluation

In order to measure the validation of our classification algorithm we used a couple of widely used performance matrices. They are defined down below:

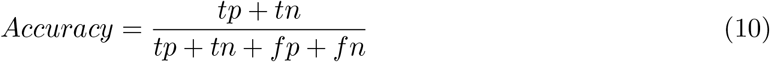

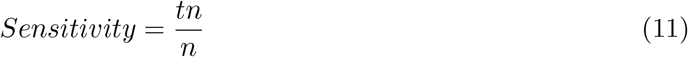

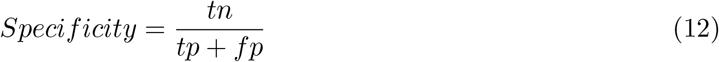

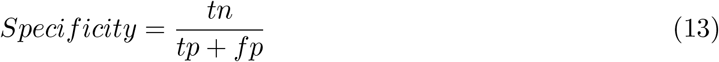

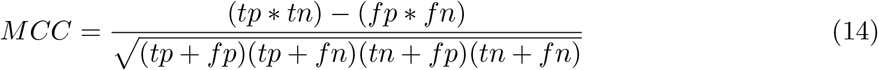

Where: *n* = number of instances of actual negative samples *p* = number of instances of actual positive samples *tp* = number of instances where positive samples are predicted correctly. *tn* = number of instances where negative samples are predicted correctly. *fp* = number of instances where negative samples are predicted incorrectly. *fn* = number of instances where positive samples are predicted incorrectly.

Accuracy,Sensitivity and Specificity have range between 0 and 1 inclusively. The perfect classifier will give value 1 and the worst one will give 0. The next one, MCC is ranged between +1 to −1, with +1 being the perfect classifier, −1 the worst and 0 is referred to as a random classifier

## 3 Experimental Analysis

The experiments were conducted using two machines, first one was equipped with Intel Core i3-8100 Processor along with one nVidia Geforce GTX 1070ti graphics card and the second machine was equipped with Intel Core i3-3100 processor and had a nVidia Geforce GTX 1050ti graphics card. The whole application was written in Python 3.6 language with using a couple of libraries including but not limited to Keras, Scikit-learn and Matplotlib.

### 3.1 Comparison with previous methods

We compared our result with 12 other methods on the benchmark train and test dataset gradually in Table 1 and Table 2. Till our proposed method DPP-PseAAC was beyond doubt the best performing classifier for training dataset with accuracy of 95.91%. Our tool, DeepDBP-ANN was a near perfect classifier for training dataset with an accuracy of 99.02%. However in order to truly determine the degree of effectiveness of a model an evaluation through a independent dataset is necessary.

**Table 1:** Comparison of DeepDBP with previous methods on PDB 1075 dataset.

| Method | Accuracy | Sensitivity | Specificity | MCC | auROC | auPR |
| --- | --- | --- | --- | --- | --- | --- |
| DNAbinder | 79.09 | 0.48 | 0.814 | 0.48 | 0.8140 | - |
| DNA-Prot | 72.55 | 0.8267 | 0.5976 | 0.44 | 0.789 | - |
| iDNA-Prot | 75.4 | 0.8381 | 0.6473 | 0.50 | 0.761 | - |
| iDNA-Prot—dis | 77.3 | 0.794 | 0.7527 | 0.54 | 0.831 | - |
| PseDNA-Pro | 76.55 | 0.79611 | 0.7363 | 0.53 | - | - |
| iDNAPro-PseAAC | 76.76 | 0.7562 | 0.7745 | 0.53 | 0.8392 | - |
| HMMBinder | 86.33 | 0.87 | 0.855 | 0.72 | 0.902 | - |
| Kmer1 + ACC | 75.23 | 0.7676 | 0.7376 | 0.50 | 0.828 | - |
| Local-DPP | 79.20 | 0.84 | 0.7450 | 0.59 | - | - |
| iDNAProt-ES | 90.18 | 0.9038 | 0.90 | 0.80 | 0.9412 | - |
| DPP-PseAAC | 95.91 | 0.941 | 0.9764 | 0.92 | 0.9884 | - |
| Grouped Feature Selection | 70.82 | 0.61 | 0.797 | 0.41 | 0.751 | 0.721 |
| Recursive Feature Selection | 71.04 | 0.62 | 0.799 | 0.43 | 0.751 | 0.724 |
| DeepDBP-ANN | **99.02** | **0.98** | **0.97** | **0.992** | **0.996** | 0.996 |
| DeepDBP-CNN | 94.32 | 0.83 | 0.75 | 0.981 | 0.986 | 0.982 |

**Table 2:** Comparison of DeepDBP with previous methods on PDB 186 validation dataset.

| Method | Accuracy | Sensitivity | Specificity | MCC | auROC | auPR |
| --- | --- | --- | --- | --- | --- | --- |
| DNAbinder | 60.80 | 0.57 | 0.645 | **0.22** | 0.216 | 0.607 |
| DNA-Prot | 61.80 | 0.68 | 0.538 | 0.24 | 0.240 | - |
| iDNA-Prot | 67.20 | 0.667 | 0.667 | 0.34 | 0.344 | - |
| iDNA-Prot—dis | 80.64 | 0.800 | 0.800 | 0.54 | 0.831 | - |
| PseDNA-Pro | 76.55 | 0.7961 | 0.7961 | 0.53 | - | - |
| iDNAPro-PseAAC | 69.89 | 0.77 | 0.624 | 0.40 | 0.8392 | 0.775 |
| HMMBinder | 69.02 | 0.61 | 0.763 | 0.39 | 0.632 | - |
| Kmer1 + ACC | 70.96 | 0.83 | 0.591 | 0.43 | 0.431 | 0.752 |
| Local-DPP | 79.00 | 0.92 | 0.656 | 0.63 | - | - |
| iDNAProt-ES | 80.64 | 0.81 | 0.800 | 0.61 | 0.843 | - |
| DPP-PseAAC | 77.42 | 0.83 | 0.709 | 0.55 | 0.798 | - |
| Grouped Feature Selection | 82.26 | 0.95 | 0.699 | 0.67 | 0.823 | 0.745 |
| Recursive Feature Selection | 76.88 | 0.77 | 0.769 | 0.55 | 0.769 | 0.696 |
| DeepDBP-ANN | 82.80 | **0.98** | **0.97** | 0.992 | **0.996** | **0.996** |
| DeepDBP-CNN | 84.31 | 0.83 | 0.75 | 0.981 | 0.986 | 0.982 |

To our knowledge, till DeepDBP-CNN, Grouped Feature Selection had been the forefront method with accuracy of 82.26%. However our second methodology, DeepDBP-CNN was able to gain even better accuracy off **84.31**%. Even though Grouped Feature Selection had a validation accuracy of 82.26%, the train accuracy wasn’t as good as the validation accuracy. Which clearly indicates the model was underfitting; which led us to believe that there were improvements to be made. So we tried DeepDBP-ANN and as predicted was able gain state-of-the-art result. But as described earlier DeepDBP-ANN had very high training accuracy compared to validation accuracy. In order to solve the overfitting problem, we tried a different methodology, a novel approach, which not only solves the overfitting problem, but also get automates the feature extraction procedure. Which can be generalized to build classifier for different datasets and even for different problems as well.

We have also performed Receiver Operating Characteristic (ROC) analysis on train and test sets for both of the methods. The graphs are shown in Figure 6. The diagonal dotted line in the middle of both the curves represents a model that is just as good as random guessing. The performance of a model is directly proportional to the area under the ROC curve. The highest value of the area can be 1.0 and in case of both the models the value of the area is greater than **0.98** on the training set and approximately **0.83** on testing set.

**Figure 5:**
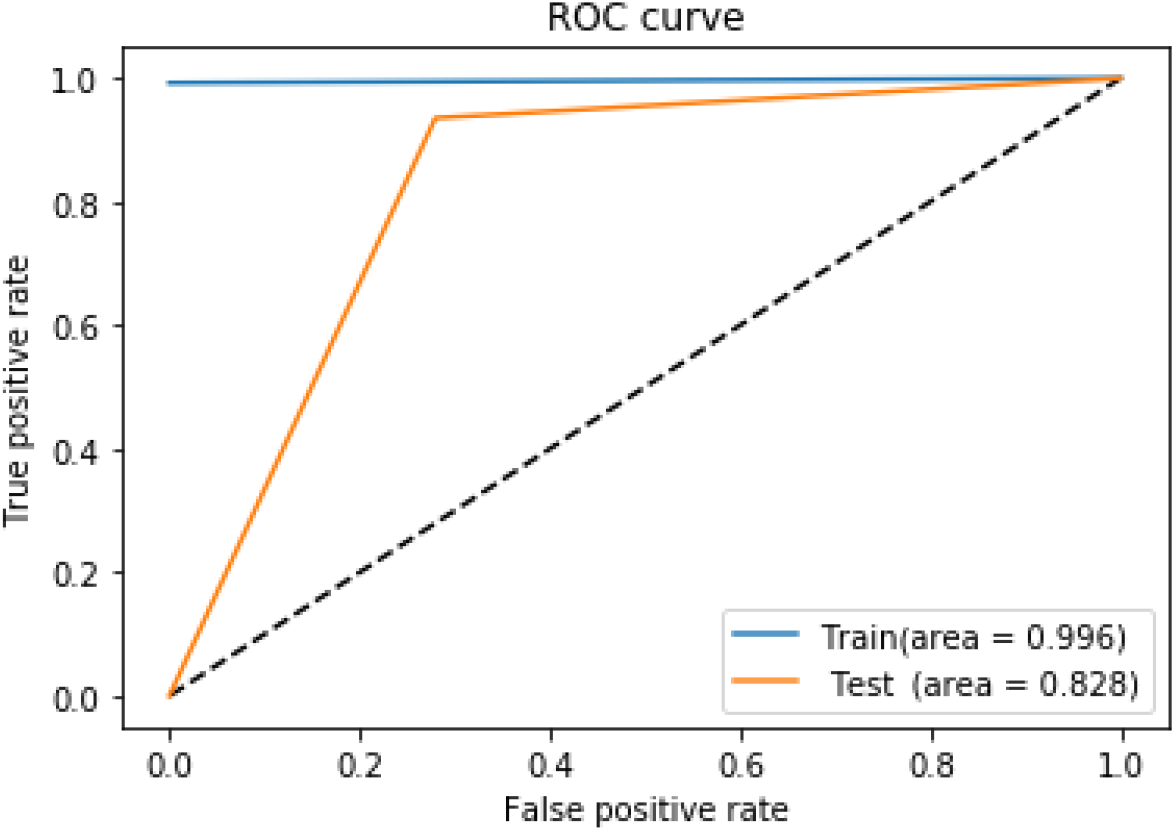
ROC curves on train and test sets for DeepDBP-ANN

**Figure 6:**
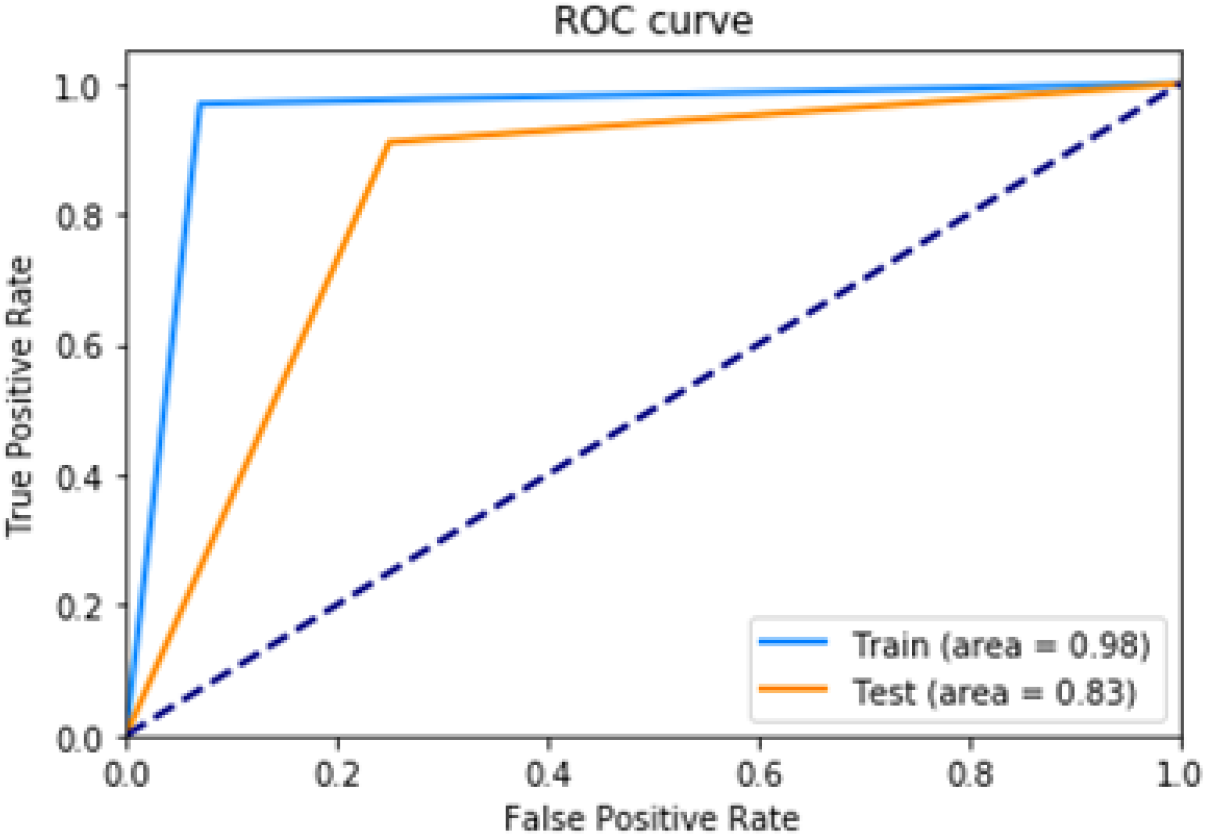
ROC curves on train and test sets for DeepDBP-CNN.

## 4 Conclusion

As described previously our deep learning model provides state of the art result with very short computation time. We tried the conventional approach of extracting features with specified algorithm and also tried the novel approach of feature extraction without any manual tweaking using deep learning techniques. While our first approach was producing state of the art result, the second approach even exceeds the result of first one. And also while the first approach is dataset specific, the second approach is more generalized, can be applied to other datasets as well and since the features are extracted by the model itself it does not require in-depth knowledge about the dataset.

